# Epigenetic profile drives accurate survival prediction in breast cancer via a multi-omics machine learning model

**DOI:** 10.1101/2025.07.31.667894

**Authors:** Guillaume Mestrallet, Alexandre Pierga, Paul Fremont

## Abstract

Accurate overall survival (OS) prediction is key for personalized treatment in breast cancer, but mutation burden alone is insufficient. To improve prognostic accuracy, we integrated genomic, transcriptomic, proteomic, epigenetic, and clinical features from 802 breast cancer patients to develop BANDOL (Breast cancer Analysis with Neoplastic Data and Omics Learning), a Random Survival Forest model. BANDOL correctly predicted survival ranking in 73% of patient pairs and outperformed mutation-based models (time-dependent AUC: 0.9–1 vs. 0.4–0.9). Immune activation signatures correlated with a longer OS after therapy, while a shorter OS was linked to TREM2⁺ myeloid cells, B cells, and leptin signaling. The transferability of the model was further explored in three independent TCGA cohorts from distinct cancer types (uterine, ovarian and lower-grade glioma), where moderate predictive performance was maintained despite biological differences between tumors. Epigenetic features were the strongest OS predictors for BANDOL. Current therapies may be combined with strategies to target the methylations of ME3, PPARG, OLIG3 and SLC25A22 genes. This study demonstrates that multi-omics integration via machine learning enhances survival prediction and reveals actionable biomarkers.

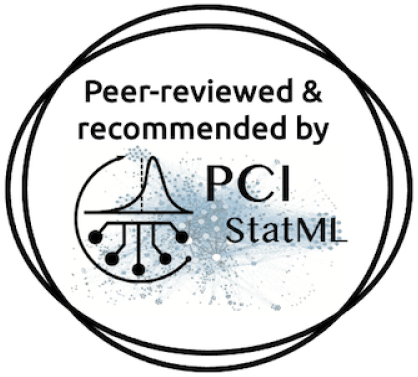

## Introduction

The accumulation of somatic mutations and the resulting increase in tumor mutational burden (TMB) have emerged as important biological features with potential clinical implications across multiple cancer types. High TMB is often associated with an increased likelihood of generating neoantigens, which may enhance tumor immunogenicity and facilitate immune-mediated tumor clearance. Consequently, a growing body of evidence suggests that elevated TMB correlates with improved responses to immune checkpoint blockade therapies and, in some cancers, with better overall survival (OS) (Dantoing et al., 2021; Mestrallet et al., 2023; Reck et al., 2019; Silva et al., 2021). OS is the length of time from either the date of diagnosis or the start of treatment for a disease, such as cancer, that patients diagnosed with the disease are still alive. Building on this observation, several studies have utilized mutational data in combination with machine learning techniques to predict patient outcomes and therapeutic responses. These approaches have demonstrated that predictive modeling using mutational features can yield clinically relevant insights and may inform personalized treatment strategies (Jee et al., 2024; Mestrallet, 2024, 2025). However, the prognostic significance of TMB is not universal. For instance, in breast cancer and other malignancies, high TMB does not reliably predict improved survival outcomes (McGrail et al., 2021; Samstein et al., 2019). This inconsistency underscores the limitations of relying solely on TMB or mutation count as prognostic biomarkers, particularly in cancers with low immunogenicity or distinct tumor microenvironments.

To address this limitation, more comprehensive models that integrate multi-dimensional tumor features are needed. A growing consensus in the field supports the incorporation of transcriptomic, proteomic, and epigenetic data, such as RNA and protein expression levels and DNA methylation patterns, alongside mutational and clinical features to improve prognostic accuracy (Menyhárt & Győrffy, 2021; Tran et al., 2025). These data layers provide complementary insights into tumor biology, capturing regulatory, functional, and structural alterations that are not reflected in mutational profiles alone. While previous studies have identified specific epigenetic or transcriptomic signatures associated with prognosis, these efforts have generally focused on isolated data types and have not employed integrative, machine learning–based approaches that could exploit the full spectrum of molecular data (Bao et al., 2019). For example, in this study (Bao et al., 2019), the authors identified an epigenetic signature associated with OS in breast cancer, and used it to design a monogram stratifying low and high-risk groups, but this approach is not as precise as machine learning approaches and does not incorporate all individual features. As such, the predictive potential of combining multi-omic features with advanced computational models remains partially underexplored, particularly in breast cancer.

In this study, we aim to fill this gap by developing a robust machine learning framework for survival prediction in breast cancer patients. We focus on breast cancer not only because of its clinical relevance and heterogeneity but also due to the absence of a clear association between TMB and patient survival, making it an ideal setting in which to evaluate the added value of integrative modeling (**Figure 1**). To this end, we leveraged data from The Cancer Genome Atlas (TCGA), which provides a rich compendium of molecular and clinical data, including tumor RNA and protein expression levels, DNA methylation profiles, mutational information, and detailed patient annotations such as OS following treatment (Ciriello et al., 2015; Koboldt et al., 2012). We trained Random Survival Forest (RSF) models, a machine learning method known for its robustness in handling high-dimensional and censored survival data, and assessed their ability to predict OS using this integrative dataset (Mestrallet, 2025; Yoo et al., 2025). To explore the generalizability of our approach beyond breast cancer, we further evaluated the model in additional TCGA cohorts representing distinct cancer types. Our findings demonstrate that integrative, multi-omic modeling substantially improves survival prediction in breast cancer patients compared to models based solely on TMB or clinical features. This approach not only advances the methodological toolkit for precision oncology but also provides new biological insights into the determinants of survival in breast cancer.

**Figure 1.**
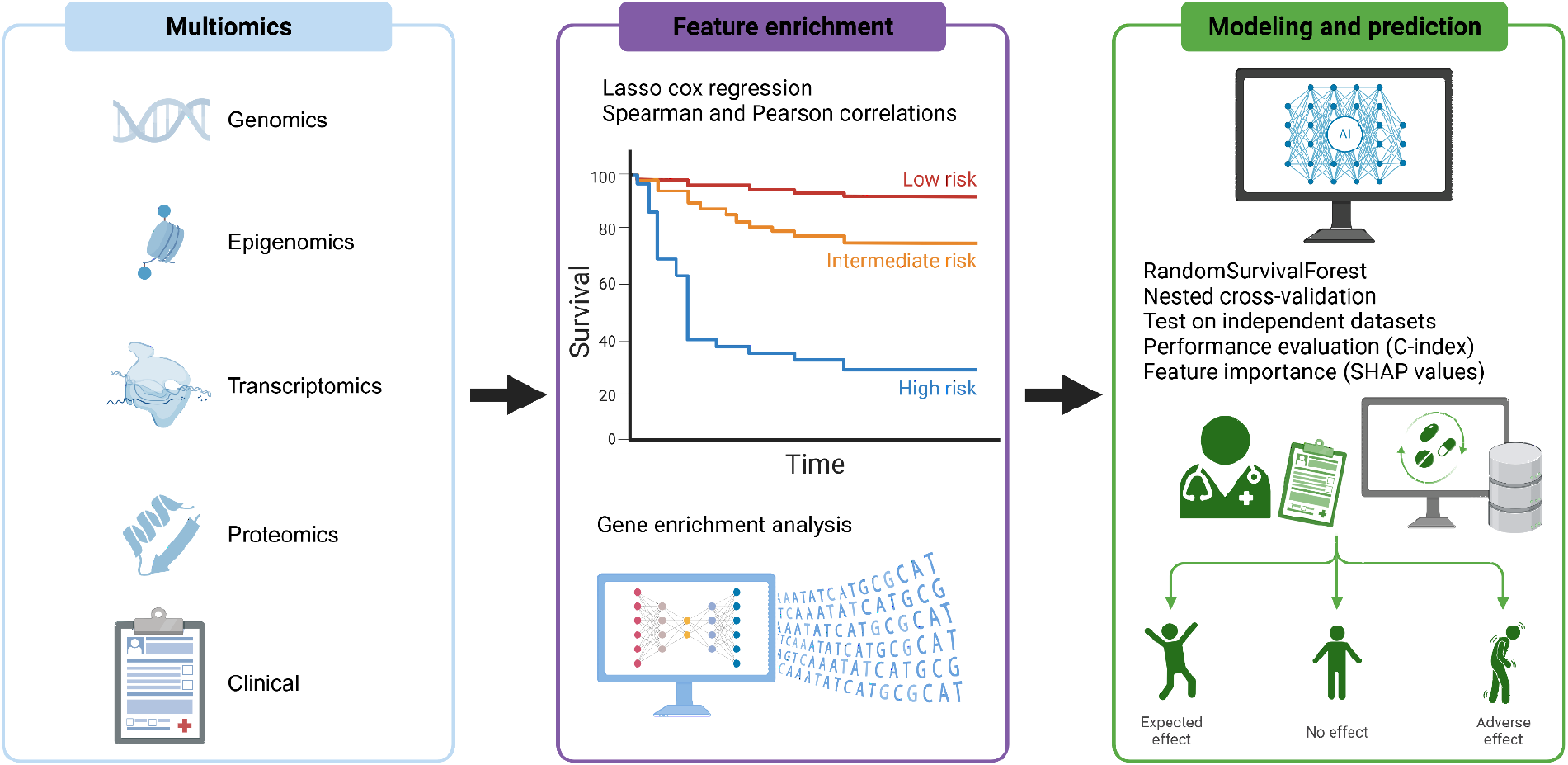
Proposed workflow to predict breast cancer patient survival using multi-omics. We analyzed multi-omic and clinical data from 802 female breast cancer patients in The Cancer Genome Atlas (TCGA), integrating genomic, transcriptomic, proteomic, epigenetic, and clinical features. Using Random Survival Forest models, we predicted overall survival based on features identified through correlation analyses, Lasso-Cox regression, and hazard ratio assessments. Model performance was evaluated via nested cross-validation and measured using the Concordance Index. Time-dependent receiver operating characteristic (ROC) curves and cumulative dynamic areas under the curve (AUC) were computed at multiple time points. Cross-cancer transferability was explored using two independent TCGA cohorts (uterine cancer, n=132; ovarian cancer, n=226; lower-grade glioma, n=189). To interpret the model and identify key prognostic drivers, we computed Shapley values for each feature.

## Results

### A limited number of mutations and clinical features correlate with OS in breast cancer

We hypothesized that the accumulation of mutations and a high TMB would not be associated with a better overall survival (OS) in breast cancer. When plotting Pearson and Spearman correlations between mutational features and OS in months, non synonymous TMB significantly negatively correlates with a longer OS, and no correlation was observed after applying the Bonferroni correction (**Figure 2 A**). Moreover, very few mutations (5 among 24496 mutations) are significantly linked to a difference in OS in months in breast cancer, and no correlation was observed after applying the Bonferroni correction (**Figure 2 B**). Of note, no mutations were associated with a shorter survival, and most of the ones significantly potentially associated with a longer survival are fusion mutations (**Figure 2 B**). Asian genetic ancestry correlates with a shorter OS after applying the Bonferroni correction, while the use of doxorubicin (blocking topoisomerase 2), paclitaxel (stopping mitosis by interfering with microtubules) or cyclophosphamide (DNA replication inhibitor) significantly correlates with a longer survival in this cohort (**Figure 2 A**). When calculating the hazard ratios and performing cox regressions, no mutations were significantly associated with a higher risk of dying in this cohort. Overall, a limited number of clinical features and no mutational features significantly correlate with OS in breast cancer.

**Figure 2.**
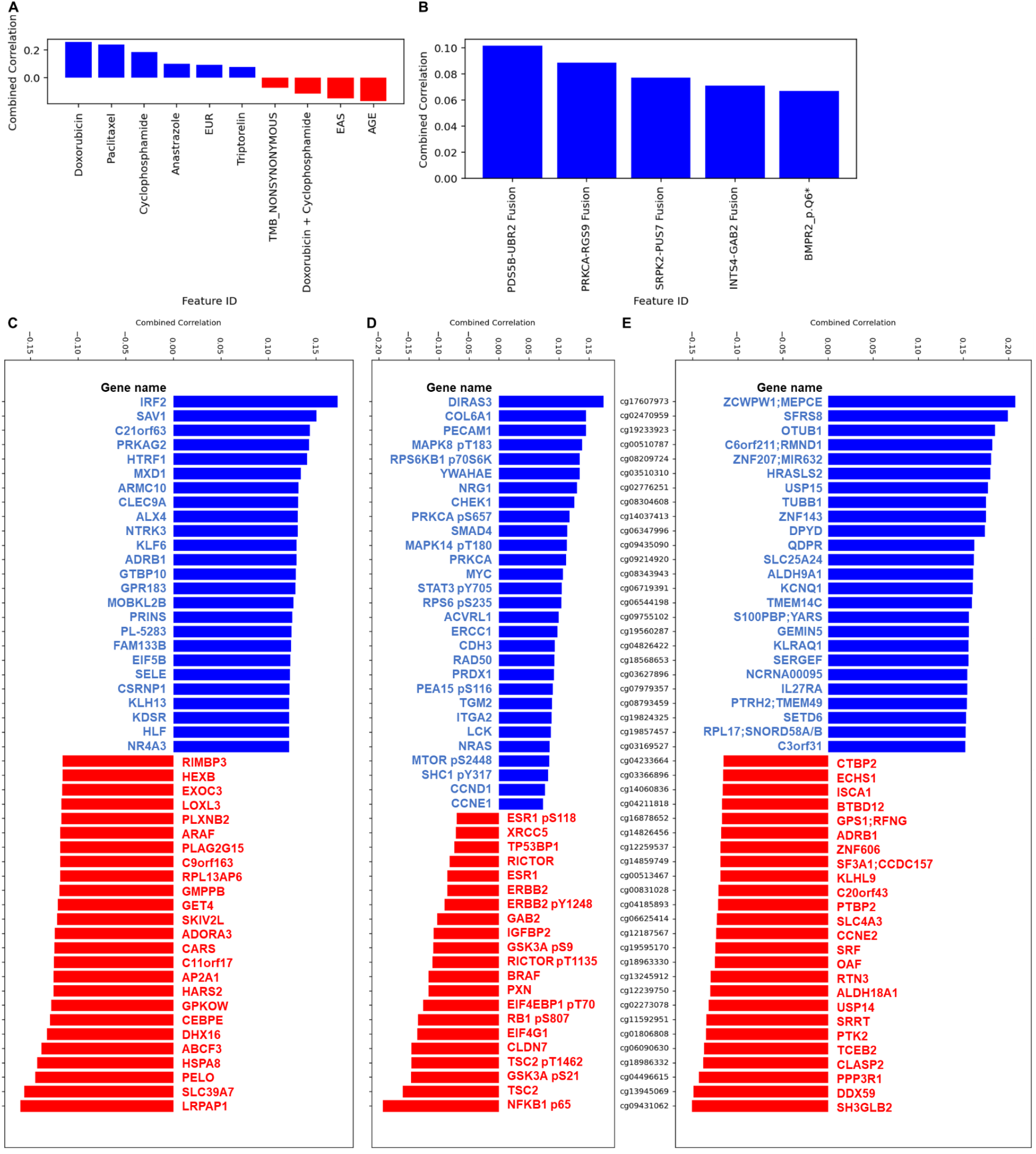
Breast cancer patient multi-omics and clinical features associated with a difference in overall survival. **A**. Breast cancer patient clinical features significantly associated with a longer or shorter overall survival (for both Pearson and Spearman correlations). Features include clinical variables, tumor mutational burden (TMB), age, genetic ancestry (EUR = europe, EAS = asia), final overall survival status (living, deceased), drug used (doxorubicin, paclitaxel, cyclophosphamide, triptorelin). The only variables plotted are the ones associated with a difference in survival using both Pearson and Spearman correlations. **B**. Breast cancer patient mutational features significantly associated with a difference in overall survival (for both Pearson and Spearman correlations). N=802 breast cancer patients. The only variables plotted are the ones associated with a difference in survival using both Pearson and Spearman correlations. **C**. Breast cancer patient mRNA transcripts (RSEM, batch normalized from Illumina HiSeq_RNASeqV2) with expression significantly associated with a longer or shorter overall survival for both Pearson and Spearman correlations. N=802 breast cancer patients. Highest and lowest 25 correlations (average between Spearman and Pearson) were plotted. The only variables plotted are the ones associated with a difference in survival using both Pearson and Spearman correlations. **D**. Breast cancer patient protein expression, measured by reverse-phase protein array (RPPA), significantly associated with a longer or shorter overall survival for both Pearson and Spearman correlations. N=802 breast cancer female patients. Highest and lowest correlations (average between Spearman and Pearson) were plotted. The only variables plotted are the ones associated with a difference in survival using both Pearson and Spearman correlations. **E**. Breast cancer patient methylations between-platform (hm27 and hm450) normalization values significantly associated with a longer or shorter overall survival for both Pearson and Spearman correlations. N=802 breast cancer patients. Highest and lowest 25 correlations (average between Spearman and Pearson) were plotted. The only variables plotted are the ones associated with a difference in survival using both Pearson and Spearman correlations. Gene enrichment and cell type signature analysis with these top/bottom genes was performed using metascape in (**Supplementary Figure 1/2**).

### An integrated genomic, transcriptomic, epigenomic, and proteomic signature predicts OS in breast cancer

We hypothesized that other multi-omic features may be associated with differences in OS and allow a better prediction in breast cancer. Importantly, we found that mRNA expression (**Figure 2 C**), protein levels (**Figure 2 D**) and DNA methylation levels (**Figure 2 E**) were significantly associated with differences in OS in breast cancer. Moreover, many of these features have a coefficient of correlation with OS between 0.1 and 0.2, while the mutation ones are all below 0.1, suggesting that mRNA, protein and DNA methylation levels are stronger predictors of OS than mutations.

In more details, we found that the mRNA expression level of IRF2, SAV1, C21orf63, PRKAG2, HTRF1, MXD1, ARMC10, CLEC9A, ALX4, NTRK3, KLF6, ADRB1, GTBP10, GPR183, MOBKL2B, PRINS, PL-5283, FAM133B, EIF5B, SELE, CSRNP1, KLH13, KDSR, HLF and NR4A3 genes significantly correlates with a longer OS (**Figure 2 C**). By applying gene set enrichment and cell type signature analysis with metascape (Y. Zhou et al., 2019), we found that this mRNA signature is close to the cytokine production (GO:0001819) and platelet-derived growth factor receptor signaling (GO:0048008) pathways, and suggests a tumor infiltration by myeloid cells (**Supplementary Figure 2)**. On the other hand, the mRNA expression level of RIMBP3, HEXB, EXOC3, LOXL3, PLXNB2, ARAF, PLAG2G15, C9orf163, RPL13AP6, GMPPB, GET4, SKIV2L, ADORA3, CARS, C11orf17, AP2A1, HARS2, GPKOW, CEBPE, DHX16, ABCF3, HSPA8, PELO, SLC39A7 and LRPAP1 genes significantly correlates with a shorter OS (**Figure 2 C**). This signature is close to mRNA metabolomic processes (GO:0016071) and translation regulation (GO:0006412) signatures (**Supplementary Figure 1)**. Overall, longer OS significantly correlated with the upregulation of myeloid, cytokine and platelet immune mRNA signatures, while shorter OS significantly correlated with the upregulation of mRNA metabolomic and translation regulation signatures.

Next, using the reverse-phase protein array (RPPA) assay, we found that the protein expression levels of DIRAS3, COL6A1, PECAM1, MAPK8, RPS6KB1, YWAHAE, NRG1, CHEK1, PRKCA, SMAD4, MAPK14, MYC, STAT3, RPS6, ACVRL1, ERCC1, CDH3, RAD50, PRDX1, PEA15, TGM2, ITGA2, LCK, NRAS, MTOR, SHC1, CCND1 and CCNE1 significantly correlate with a longer OS (**Figure 2 D**). By applying gene set enrichment and cell type signature analysis with metascape, we found that this proteomic signature is close to the IL-2, and type 1 IFN pathways and suggests a tumor infiltration by naive T cells (**Supplementary Figure 2)**. On the other hand, the protein expression levels of ESR1, XRCC5, TP53BP1, RICTOR, ESR1, ERBB2, ERBB2, GAB2, IGFBP2, GSK3A, RICTOR, BRAF, PXN, EIF4EBP1, RB1, EIF4G1, CLDN7, TSC2, GSK3A, TSC2 and NFKB1 significantly correlate with a shorter OS (**Figure 2 D**). This signature is close to the leptin and multiple cancer immunosuppressive signatures (**Supplementary Figure 1)**. Overall, longer OS significantly correlated with the upregulation of IL-2 and type 1IFN pathway proteomic signatures, while shorter OS significantly correlated with the upregulation of leptin and multiple cancer immunosuppressive proteomic signatures.

Finally, we found that the DNA methylation level of ZCWPW1, MEPCE, SFRS8, OTUB1, C6orf211, RMND1, ZNF207, MIR632, HRASLS2, USP15, TUBB1, ZNF143, DPYD, QDPR, SLC25A24, ALDH9A1, KCNQ1, TMEM14C, S100PBP, YARS, GEMIN5, KLRAQ1, SERGEF, NCRNA00095, IL27RA, PTRH2, TMEM49, SETD6, RPL17, SNORD58A, SNORD58B and C3orf31 genes significantly correlates with a longer OS (**Figure 1 E**). On the other hand, the DNA methylation level of CTBP2, ECHS1, ISCA1, BTBD12, GPS1, RFNG, ADRB1, ZNF606, SF3A1, CCDC157, KLHL9, C20orf43, PTBP2, SLC4A3, CCNE2, SRF, OAF, RTN3, ALDH18A1, USP14, SRRT, PTK2, TCEB2, CLASP2, PPP3R1, DDX59 and SH3GLB2 genes significantly correlates with a shorter OS (**Figure 2 E**). Thus, differences in OS are significantly associated with multiple DNA methylation signatures.

### Treatment response is linked to innate and adaptive immune activation, while resistance correlates with immunosuppressive leptin signatures

To better characterize the pathways associated with a difference in survival, we performed a fully integrated gene enrichment and cell type signature analysis with these genomic, transcriptomic and proteomic signatures combined using metascape (**Figure 3 A**). From an immunological perspective, we observed that a longer OS was associated with the presence of naive T cells and myeloid cells (monocytes and dendritic cells), IL-2 and IFN type 1 signaling pathways, while a shorter OS was associated with TREM2 immunosuppressive myeloid cell (Binnewies et al., 2021; Mestrallet et al., 2026) and B cell signatures, and the leptin pathway known to be key in immune regulation (Park & Ahima, 2014) (**Figure 3 A**).

**Figure 3.**
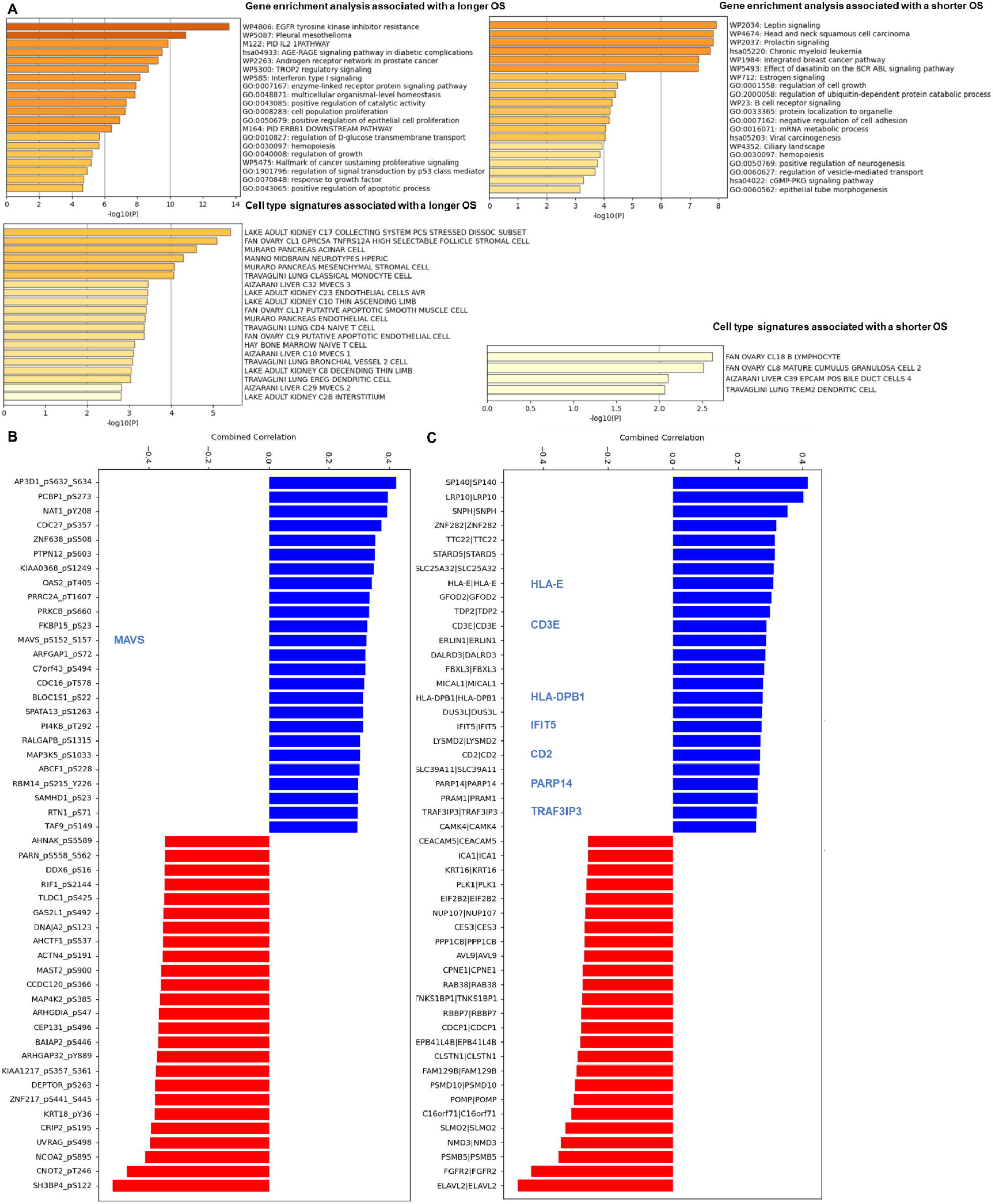
Treatment response is linked to innate and adaptive immune activation, while resistance correlates with immunosuppressive leptin signatures. **A**. Gene enrichment and cell type signature analysis significantly associated with a difference in survival. Gene enrichment and cell type signature analysis with these top/bottom genes (from the RNA dataset of Figure 1 C, from the methylation dataset of Figure 1 D and from the protein dataset of Figure 1 E) was performed using metascape (Y. Zhou et al., 2019). **B**. Breast cancer patient protein expression by mass spectrometry and their phosphorylation levels significantly associated with a difference in survival. **B/C**. Breast cancer patient protein expression (**B**) and their level of phosphorylation (**C**), directly measured on 56 patients, significantly associated with a longer or shorter overall survival for both Pearson and Spearman correlations. N=56 female breast cancer patients. Highest and lowest correlations (average between Spearman and Pearson) were plotted. The only variables plotted are the ones associated with a difference in survival using both Pearson and Spearman correlations.

To investigate it further, we looked at the direct protein expression using mass spectrometry and not RPPA as we did before, and their level of phosphorylation in the tumors of the 56 patients that were profiled in the cohort (**Figure 3 B/C**). We confirmed that longer survival was associated with antigen presentation to T cells (HLA-DPB1 expression), T cell infiltration (CD3E expression) and NK cell infiltration (HLA-E and CD2 expression) (**Figure 3 B/C**). Moreover, MAVS phosphorylation and downstream Interferon Stimulated Genes (ISGS) pathway (MAVS, TRAF3IP3, IFIT3, IFIT5, PARP14, FADD, CAMK2D, RFX5, GBP4, PSMB8 and BST2) activation significantly correlate with survival in breast cancer. The MAVS pathway, essential for the antiviral immune response, functions as a central hub that connects virus recognition to innate immunity (Ren et al., 2020). TRAF3IP3 is a protein that interacts with TRAF3 and is involved in downstream MAVS signaling, mediating the recruitment of TRAF3 to MAVS for effective antiviral responses (Zhu et al., 2019). IFIT5 plays a significant role in recognizing viral RNA and participating in downstream signaling that enhances antiviral mechanisms (Zhang et al., 2013). Finally, PARP14 regulates immune signaling pathways, including those mediated by MAVS, influencing antiviral responses (Du et al., 2023; Parthasarathy & Fehr, 2022). Overall, response to therapy significantly correlated with the activation of both innate and adaptive immunity, while resistance significantly correlated with TREM2+ myeloid, B cell and immunosuppressive leptin signatures.

### Epigenetic profile drives accurate survival prediction in breast cancer via a multi-omics machine learning model

We hypothesized that we may leverage machine learning algorithms trained on the multi-omics features that we identified in **Figure 2** as associated with a higher or lower risk to better predict cancer patient outcome. First, we investigated more deeply which multi-omics breast cancer patient features are significantly associated with a difference in survival in this cohort of 802 patients using a Lasso cox regression. This test takes into account not only the OS time component but also the event/status of the patient at the end point (deceased/living). Lower risks were observed in presence of higher ESRRG, IRF2 and MAP2K6 RNA levels, and cg04515986 (FTHL17 gene), cg06531741 (RIN2 gene) and cg17016000 (HTR3B gene) methylation levels, and the use of Cyclophosphamide (**Figure 4 A**). On the contrary, increased risks were observed for older patients, and patients with higher PELO and MAFA RNA levels, phosphorylated NFKB1 protein levels and cg09645888 (ME3 gene), cg03889226 (OLIG3 gene), cg04632671 (PPARG gene) and cg10970251 (SLC25A22 gene) methylation levels (**Figure 4 A**).

**Figure 4.**
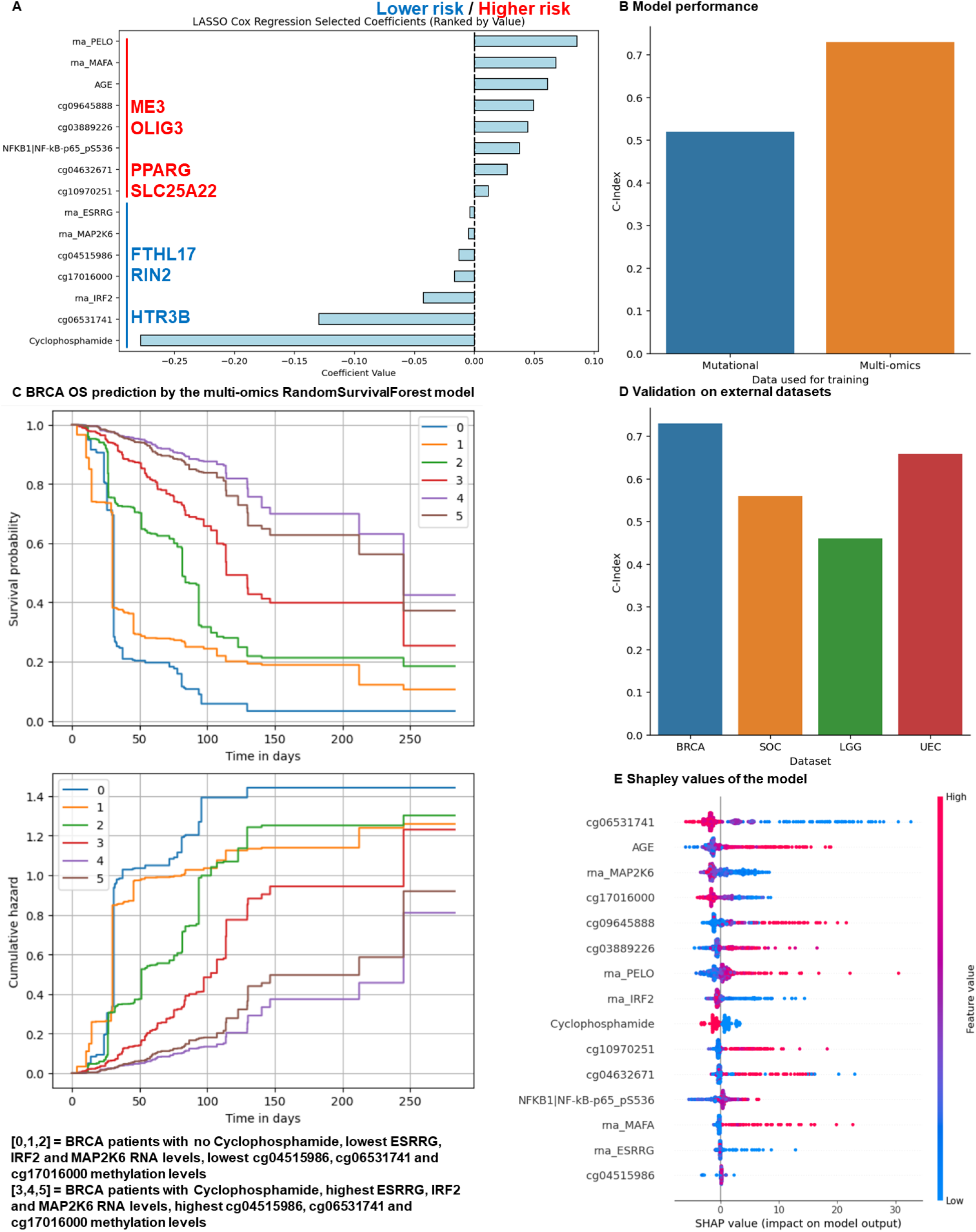
Epigenetic profile drives accurate survival prediction in breast cancer via a multi-omics machine learning model. **A**. Cancer patient features associated with an increased or decreased hazard ratio (Lasso cox regression). N=802 breast cancer female patients. **B**. Prediction of overall survival of cancer patients based on their methylation, RNA, protein and clinical features identified in panel 3 A. A RandomSurvivalForest model was used with nested cross validation. Survival was predicted using the Kaplan-Meier estimator. Model performance (C-Index) according to the dataset used for training (multi-omics vs mutational data). **C**. Prediction of OS and the cumulative risk (estimating the probability of experiencing the event (e.g., death) by a certain time) based on ESRRG, IRF2 and MAP2K6 RNA levels, and cg04515986, cg06531741 and cg17016000 methylation levels, and the use of Cyclophosphamide. **D**. The model was tested on 2 other independent datasets (132 uterine cancer (UEC), 226 ovarian cancer (SOC), 189 lower grade glioma (LGG) female patients). **E**. Features predicting response by the model (Shapley values).

We used a Random Survival Forest model trained on these features (**Table 1**) with nested cross-validation and parameter optimization to predict the OS status (0 for living, 1 for deceased) and OS in months. The outer loop consisted of a 5-fold cross-validation to estimate the generalization performance of the model, while the inner loop performed 5-fold cross-validation on the training data to select optimal hyperparameters via grid search (**Supplementary Table 1**). Model performance was evaluated using the concordance index (C-index) computed on held-out test folds in the outer loop. To obtain a final model for downstream application, we retrained the RSF model on the entire dataset using the hyperparameter set that achieved the best average cross-validation performance in the inner loop. We obtained a c-Index of 0.73 for the model trained on multi-omics data vs 0.52 for the one trained only on mutation data, meaning that the multi-omics model was more efficient to predict OS (**Figure 4 B**). It indicates that in 73% of cases, the model correctly predicts that a patient with a shorter predicted survival time indeed has a shorter observed survival time compared to another patient, and globally outperformed models trained on mutation profile.

**Table 1.** Features selected for BANDOL training.

| Feature type |  |  |  |  |  |  |  |
| --- | --- | --- | --- | --- | --- | --- | --- |
| Methylation | SLC25A22 | PPARG | OLIG3 | FTHL17 | ME3 | RIN2 | HTR3B |
| mRNA | PELO | MAFA | ESRRG | MAP2K6 | IRF2 |  |  |
| Clinical | Age | Cyclophosphamide |  |  |  |  |  |
| Protein | NFKB1 |  |  |  |  |  |  |

As an example, we plotted the prediction of OS and the cumulative risk (estimating the probability of experiencing the event (e.g., death) by a certain time) based on ESRRG, IRF2 and MAP2K6 RNA levels, and cg04515986, cg06531741 and cg17016000 methylation levels, and the use of Cyclophosphamide for several patients (**Figure 4 C**). On each graph, patients 0, 1 and 2 have the lowest levels of the feature of interest, while patients 3, 4 and 5 have the highest levels. It shows that, as expected, patients with lower levels of these features have an increased predicted risk compared to patients with higher levels of these features.

Next, we explored the transferability of the breast cancer-derived model to two independent TCGA cohorts representing distinct cancer types, and obtained a c-Index of 0.66 for 132 uterine cancer (UEC) patients, a c-Index of 0.56 for 226 ovarian cancer (SOC) patients and a c-Index of 0.47 for 189 lower grade glioma (LGG) patients (**Figure 4 D**). This multi-omics model outperformed the model trained on uterine and ovarian cancer mutation profiles but not glioma mutation profiles (**Table 2**). These results suggest that some prognostic signals captured by the model may be shared across cancer types, although predictive performance decreases relative to breast cancer, likely reflecting biological differences between tumor entities.

**Table 2.** Model performance (C-Index) depending on training features.

| Cancer type\Features | Mutational only C-Index | Multi-Omics C-Index |
| --- | --- | --- |
| **BRCA** | **0.52** | **0.73** |
| **UEC** | **0.55** | **0.66** |
| **LGG** | **0.50** | **0.46** |
| **SOC** | **0.51** | **0.56** |

We further investigated which features drive the prediction of survival in our model by calculating the Shapley values (**Figure 4 E**). We observed that the features with the highest importance to drive the model performance are the epigenetic ones. In more details cg06531741 (HTR3B), cg17016000 (RIN2), cg09645888 (ME3) and cg03889226 (OLIG3) methylations predict overall survival by our machine learning model (**Figure 4 E**). Altogether, training machine-learning models on clinical and multi-dimensional genomic features could better predict overall survival, and DNA methylations lead these predictions.

To address the possibility that Lasso-Cox may favor variables predictive in a linear model rather than a nonlinear Random Survival Forest, we repeated feature selection using Random Survival Forest permutation importance for feature selection instead of Lasso-Cox (**Supplementary Figure 3 A**). We observed 33% of overlaps between the 2 methods, with 5 features selected by the 2 approaches (AGE, Cyclophosphamide, Cg09645888 (ME3), Cg17016000 (RIN2), RNA PELO). Models trained on Lasso-Cox selected features achieved a similar C-index (0.73) than models trained on RSF-selected features (0.75) (**Supplementary Figure 3 B**), indicating that the predictive signal is robust to the feature-selection strategy.

To evaluate the prognostic performance of our models, we generated time-dependent receiver operating characteristic (ROC) curves and computed the cumulative dynamic area under the curve (AUC) at multiple time points (**Figure 5**). The time-dependent AUC reflects the model’s ability to discriminate between individuals who experience an event (e.g., death) by a given time versus those who do not, accounting for censoring. An AUC close to 1 indicates strong predictive performance, while 0.5 corresponds to random chance. Additionally, we plotted the Kaplan–Meier survival curve based on observed outcomes in the test cohorts, alongside the average predicted survival curve derived from the model. This comparison provides a visual assessment of how closely the model’s predicted survival aligns with actual patient outcomes over time. In accordance with the c-Index results, the multi-omics model outperformed the mutational model for test BRCA data, with time-dependent AUC between 0.9 and 1 (**Figure 5 A**) vs 0.4 and 0.9 (**Figure 5 B**). Overall, our multi-omics model outperformed the mutational model performances to predict OS.

**Figure 5.**
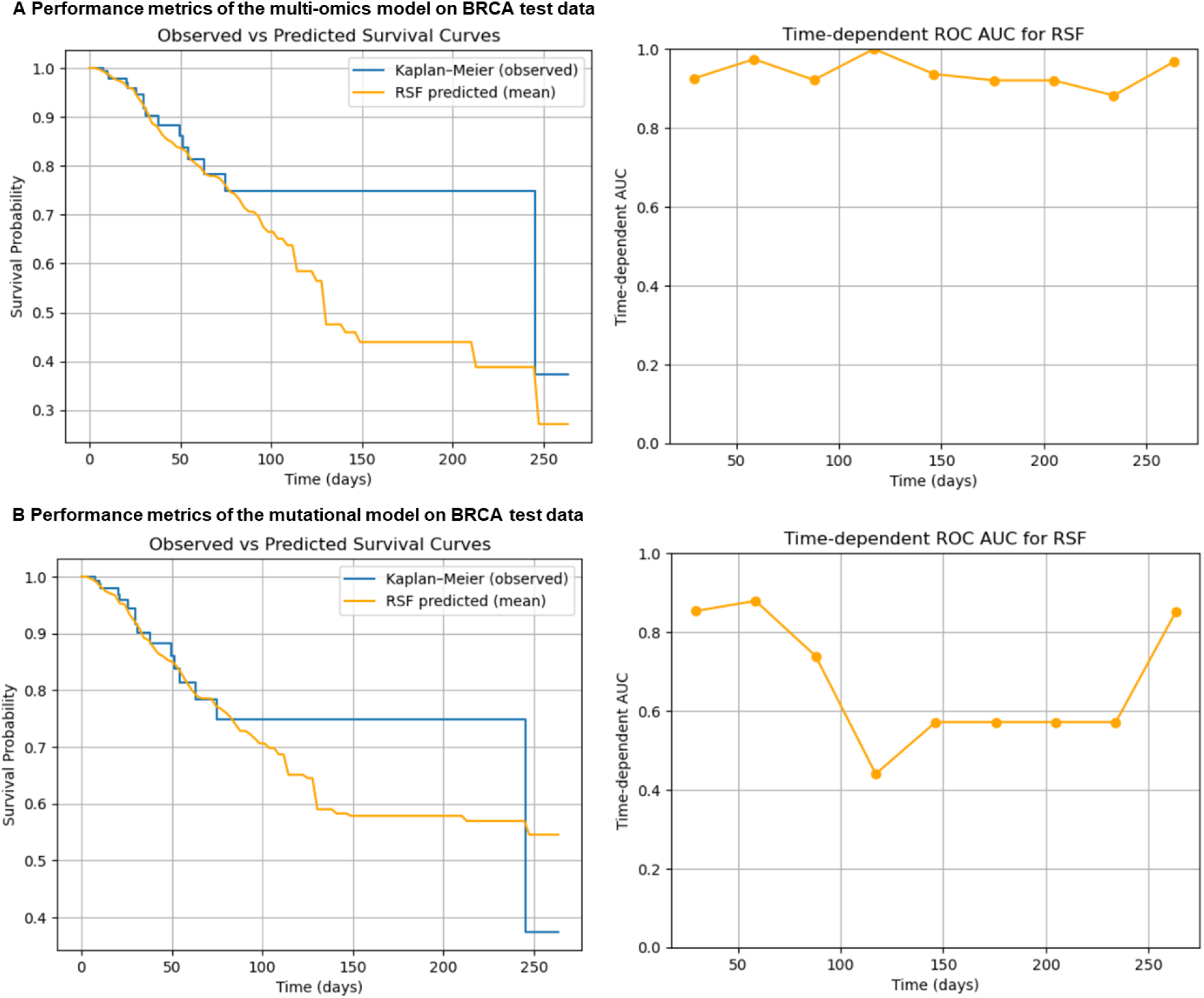
Performance metrics of the multi-omics and mutational models on the test data. **A**. Performance metrics of the multi-omics model on BRCA test data. **B**. Performance metrics of the mutational model on BRCA test data. Time-dependent receiver operating characteristic (ROC) curves and cumulative dynamic areas under the curve (AUC) were computed at multiple time points. Kaplan–Meier survival curves based on observed outcomes in the test cohorts were also plotted alongside the average predicted survival curves derived from the models.

## Discussion

Accurate prediction of overall survival (OS) in breast cancer patients is crucial for personalized treatment strategies. However, mutation profiling and a high TMB do not predict a better OS in all cancers. To better predict breast cancer patient outcomes, we considered approaches integrating clinical features with multi-dimensional genomic (mutations), epigenetic (DNA methylations) and proteomic features from patient tumors.

We observed that lower risks were observed in presence of higher ESRRG, IRF2 and MAP2K6 RNA levels. In several cancers, Interferon regulatory factor 2 (IRF2) downregulation was associated with worst outcomes by limiting MHC class I antigen presentation and increased PD-L1 expression (Kriegsman et al., 2019). However, in esophageal cancers, IRF2 promotes resistance to IFN-II and immune escape by inhibiting transcription of the IFNγR (Lukhele et al., 2022). Similarly, estrogen-related receptor gamma (ESRRG) expression was associated with better outcomes in several cancers such as gastric cancer (GC) (Kang et al., 2018), but not in esophageal squamous cell carcinoma (ESCC) (Wang et al., 2023). This gene encodes a member of the estrogen receptor-related receptor (ESRR) family. Finally, mitogen-activated protein kinase kinase 6 (MAP2K6) expression is downregulated in non-responder colorectal cancer patients of oxaliplatin therapy (Rasmussen et al., 2016). MAP2K6 phosphorylates and activates p38 MAP kinase in response to inflammatory cytokines or environmental stress, and regulates stress induced cell cycle arrest, transcription activation and apoptosis. Lower risks were also observed in presence of higher cg04515986 (FTHL17 gene), cg06531741 (RIN2 gene) and cg17016000 (HTR3B gene) methylation levels. In breast cancer, the expression of 5-hydroxytryptamine receptor (HTR2A/2B) was positively and significantly correlated with the infiltration of immune cells such as CD8+ T cells and macrophages (Zhan et al., 2023). It was also shown that NAA20 promotes the growth and invasion capability of triple-negative breast cancer cells by recruiting Ras and Rab interactor 2 (RIN2) to inhibit Rab5A-mediated activation of EGFR signaling (Qiao et al., 2023). Moreover, ferritin heavy polypeptide-like 17 (FTHL17) gene is a cancer/testis antigen gene that is overexpressed in cancer cells and in testis (Aoki et al., 2017). Thus, epigenetic regulation of these genes may disrupt the underlying immune escape mechanisms, and response to therapy significantly correlated with the activation of both innate and adaptive immunity.

On the other hand, increased risks were observed for older patients, and patients with higher PELO and MAFA RNA levels, as well as for patients with elevated phosphorylation of NFKB1. Interestingly, PELO was recently identified as a promising drug target for MSI-H patients with deleterious TTC37 mutations or with biallelic 9p21.3 deletions involving FOCAD (Borck et al., 2025). MAFA is a MAF transcription factor and acts as a transactivator of insulin and several genes involved in glucose-stimulated insulin secretion (Fottner et al., 2022). MAFA is up-regulated in 50% of human multiple myelomas and 60% of angioimmunoblastic T cell lymphomas (Iacovazzo et al., 2018). In breast cancer, NFKB1, which encodes the p50 subunit of the NF-κB protein complex, is often dysregulated and phosphorylated, leading to the activation of the NF-κB signaling pathway (Pavitra et al., 2023). This activation is frequently associated with tumor growth, metastasis, and treatment resistance. Higher risks were also observed in patients with higher cg09645888 (ME3 gene), cg03889226 (OLIG3 gene), cg04632671 (PPARG gene) and cg10970251 (SLC25A22 gene) methylation levels. It was shown that targeting Malic Enzyme 3 (ME3) may be a strategy to treat pancreatic ductal adenocarcinoma (PDAC), as overexpression of ME3 promotes pancreatic tumor proliferation, invasiveness and metastasis (Grell et al., 2022). ME3 encodes a mitochondrial NADP(+)-dependent enzyme that catalyzes the oxidative decarboxylation of malate to pyruvate, a crucial step in energy production and biosynthesis. OLIG3 is a transcription factor that regulates early cerebellar development, cell proliferation and differentiation (Lowenstein et al., 2021), and other studies have established functional roles of OLIG1 and OLIG2 in brain cancer (Szu et al., 2023). Peroxisome proliferator-activated receptor gamma (PPARG) is a ligand-dependent transcription factor expressed that regulates tumor development, progression, and metastasis in breast cancer (Augimeri et al., 2020), and it also promotes mitochondrial biogenesis(Miglio et al., 2009). It was shown that epigenetic derepression converts PPARG into a druggable target in triple-negative and endocrine-resistant breast cancers (Loo et al., 2021). It was also shown that targeting of SLC25A22, a mitochondrial glutamate carrier (Shin et al., 2023), boosts the immunotherapeutic response in KRAS-mutant colorectal cancer by suppressing CXCL1 production and impairing MDSC recruitment (Q. Zhou et al., 2023). Thus, targeting these molecules regulating mitochondrial activity in high risk patients may improve their OS (Mukherjee et al., 2023).

Altogether, response to therapy significantly correlated with the activation of both innate and adaptive immunity in the tumor, while resistance significantly correlated with TREM2+ myeloid, B cell and immunosuppressive leptin signatures. Leveraging on this multi-omics signature, we developed BANDOL, a RandomSurvivalForest model that estimates OS of 802 breast cancer patients following training on these clinical and multi-omics data associated with a difference in OS. In 75% of cases, BANDOL correctly predicts that a patient with a shorter predicted survival time indeed has a shorter observed survival time compared to another patient. The model retained moderate predictive performance when applied to distinct cancer types, suggesting partial transferability of prognostic signals, although cancer-specific biological factors likely limit performance outside the breast cancer setting. Thus, BANDOL can compute a robust ‘survival probability’ or ‘risk’ for each patient at a given time after it enters treatment. Overall, integrating not only genomic, but also transcriptomic, proteomic, and most importantly epigenetic profiles using machine learning is promising to better predict cancer patient OS. We showed that epigenetic features are the most important ones to drive the prediction by our model. Current therapies may be combined with strategies to target ME3, PPARG, OLIG3 and SLC25A22 gene methylations, as well as strategies to dephosphorylate NFKB1 and target MAFA and PELO.

Several limitations should be considered. First, initial feature selection was performed using Lasso-Cox regression prior to Random Survival Forest training. While this strategy efficiently reduces dimensionality in high-dimensional datasets, variables selected by a linear model are not necessarily optimal for a nonlinear model. However, Models trained on Lasso-Cox selected features achieved a similar C-index (0.73) than models trained on RSF-selected features (0.75), indicating that the predictive signal is robust to the feature-selection strategy. Future studies could evaluate alternative approaches such as Elastic Net-Cox regression (Greenwood et al., 2020), which better accommodates correlated predictors, or consensus features nested cross-validation (Parvandeh et al., 2020). Second, although our study demonstrates the predictive value of integrating multi-omics features, clinical-only models trained on substantially larger cohorts have achieved comparable predictive performance (Li et al., 2024). Finally, the external cohorts used in this study corresponded to different cancer types rather than independent breast cancer datasets. Consequently, these analyses should be interpreted as an assessment of cross-cancer transferability rather than a formal external validation of breast cancer survival prediction.

## Methods

### Patient cohorts

We analyzed data from The Cancer Genome Atlas (TCGA) Breast Invasive Carcinoma (BRCA) cohort (data release 43, GRCh38/hg38), a publicly available, large-scale dataset that provides multi-omic and clinical information for breast cancer patients. This cohort comprises over 1,000 primary breast cancer samples from female patients and is uniquely suited for integrative analyses aimed at uncovering molecular determinants of clinical outcomes.

The TCGA BRCA dataset includes several molecular data types:

- Genomic data: Whole-exome sequencing (WES) data were used to identify somatic single nucleotide variants (SNVs), small insertions and deletions (indels), and copy number alterations (CNAs), allowing for comprehensive mutational profiling of tumors.
- Transcriptomic data: RNA sequencing (RNA-seq) and microarray data were used to quantify gene expression levels across the transcriptome.
- Epigenomic data: Genome-wide DNA methylation profiles were obtained using Illumina Infinium HumanMethylation arrays, providing insight into regulatory changes associated with gene expression and silencing.
- Proteomic data: Reverse-phase protein array (RPPA) data measured relative protein abundance and post-translational modifications across signaling and metabolic pathways. Mass spectrometry was also performed on 56 patients.
- Clinical data: Curated clinical annotations include tumor stage, histologic grade, hormone receptor status (estrogen receptor [ER], progesterone receptor [PR]), HER2 status, treatment regimens, and patient outcomes, including overall survival (OS), disease-free survival (DFS), and response to therapy. OS is the length of time from either the date of diagnosis or the start of treatment for a disease, such as cancer, that patients diagnosed with the disease are still alive.

To test the generalizability of our model and evaluate its robustness across cancer types, we included three independent validation cohorts also obtained from TCGA (132 uterine cancer (UEC), 226 ovarian cancer (SOC) and 189 lower grade glioma (LGG) patients).

All patients included in this study were biologically female, and informed consent and ethical approvals were handled by the original TCGA consortium.

### Statistical analyses

To identify tumor and clinical features associated with patient survival, we applied multiple statistical approaches. First, Pearson and Spearman correlation analyses were used to examine linear and rank-based associations, respectively, between individual features and overall survival time. This allowed us to assess both non-linear (Spearman) and linear (Pearson) relationships between continuous molecular measurements (e.g., gene expression, methylation levels, protein abundance) and survival duration. Bonferroni correction was also applied for multiple comparisons.

Next, we performed Lasso penalized Cox proportional hazards regression to identify the most predictive features while controlling for multicollinearity and reducing overfitting in high-dimensional datasets (Jardillier et al., 2022). This is a statistical method that combines the Cox proportional hazards model with Lasso regularization, enabling variable selection and shrinkage in survival analysis. This approach is particularly useful when dealing with high-dimensional data, where the number of potential predictor variables exceeds the number of observations. Lasso regularization was used to shrink less informative feature coefficients toward zero and reduce dimensionality across multiple data modalities. We note that Lasso-based feature selection may be unstable in the presence of highly correlated predictors, and alternative approaches such as Elastic Net regularization may offer advantages in some multi-omics settings (Greenwood et al., 2020).

We also calculated hazard ratios (HRs) and corresponding 95% confidence intervals for individual features to estimate their relative impact on the risk of death. A HR measures how often a particular event happens in one group compared to how often it happens in another group, over time (Barraclough et al., 2011). A hazard ratio of one means that there is no difference in survival between the two groups. A hazard ratio of greater than one or less than one means that survival was better in one of the groups. These analyses included a range of data types—clinical variables (e.g., age, tumor stage), mutational features (e.g., TMB, driver mutations), transcriptomic and proteomic levels, and epigenetic marks (e.g., DNA methylation levels). Statistical analyses were performed using Python libraries (scipy, statsmodels, lifelines and sksurv). The final curated dataset included N = 802 breast cancer patients with complete clinical and molecular data.

### Machine-learning models for survival prediction

To develop predictive models of overall survival (OS) in breast cancer, we implemented machine learning workflows using features identified in the previous statistical analyses (**Table 1**). Our goal was to construct models that could capture complex, nonlinear relationships between multi-omic tumor features and patient outcomes.

We employed the Random Survival Forest (RSF) algorithm (Yoo et al., 2025), an ensemble tree-based method well-suited for high-dimensional survival data with right-censoring. RSF extends the standard random forest algorithm to survival analysis by growing survival trees using log-rank splitting criteria. This method does not assume proportional hazards and is robust to multicollinearity and missing data.

The input features (**Table 1**) for the model included:

- Clinical variables: age at diagnosis, treatment information
- Mutational features: TMB, specific driver gene mutations
- Transcriptomic features: normalized RNA expression levels
- Proteomic features: RPPA-based protein levels
- Epigenetic features: normalized methylation beta-values for selected CpG sites

For each model configuration, feature selection was performed independently within the training data of each outer cross-validation fold using Lasso-Cox regression. The regularization parameter was selected by cross-validation within the training set. This procedure was repeated separately for the mutational-only model and the multi-omics model to avoid information leakage from the test folds. We selected features using Lasso-Cox regression prior to Random Survival Forest training to reduce dimensionality and mitigate overfitting in the high-dimensional multi-omics dataset.

As an alternative feature selection strategy, we evaluated Random Survival Forest (RSF) feature importance. First, an RSF model was trained using all available candidate variables. Feature importance were then computed for each feature using the fitted RSF model predictions. Features were ranked according to their mean absolute importance across all patients, reflecting their overall contribution to model predictions. In more detail, we trained 4,000 models, each using 40 variables. Variable importance was assessed using the model’s feature importance metric. Variables were then ranked primarily by how frequently they appeared in the top 5 most important features across models, and secondarily by their mean importance score to break ties. The top 15 ranked features were subsequently retained and used to train a new RSF model, whose performance was evaluated using the same cross-validation framework as the primary analysis. This approach was compared with the Lasso-Cox feature selection strategy used in BANDOL. We observed 33% of overlaps between the 2 methods, with 5 features selected by the 2 approaches (AGE, Cyclophosphamide, Cg09645888 (ME3), Cg17016000 (RIN2), RNA PELO) (Supplementary Figure 3 A). Models trained on Lasso-Cox selected features achieved a similar C-index (0.73) than models trained on RSF-selected features (0.75) (Supplementary Figure 3 B), indicating that the predictive signal is robust to the feature-selection strategy.

To assess the predictive performance of clinical and molecular features on overall survival, we implemented a nested cross-validation approach using the RSF model (Cawley & Talbot, 2010; Stone, 1974). The outer loop consisted of a 5-fold cross-validation to estimate the generalization performance of the model, while the inner loop performed 5-fold cross-validation on the training data to select optimal hyperparameters via grid search. The hyperparameter search space and optimal parameter values identified during nested cross-validation are reported in Supplementary Table 1. Model performance was evaluated using the concordance index (C-index) (Yoo et al., 2025) computed on held-out test folds in the outer loop, a standard metric for survival prediction that evaluates the proportion of correctly ranked survival times. A C-index of 0.5 corresponds to random prediction, while a value of 1.0 indicates perfect prediction. To obtain a final model for downstream application, we retrained the RSF model on the entire dataset using the hyperparameter set that achieved the best average cross-validation performance in the inner loop. All analyses were performed using Python (scikit-learn and scikit-survival packages), with joblib for parallelization. Then we tested the final model on two external cohorts (UEC, SOC, LGG).

To interpret the model and identify key molecular and clinical drivers of survival, we computed Shapley Additive Explanation (SHAP) values (Mestrallet, 2025; Yoo et al., 2025), which quantify the contribution of each feature to the model’s prediction for each individual. SHAP values offer a unified approach to feature attribution and facilitate biological interpretation by ranking features based on their average impact on model outputs.

To evaluate the prognostic performance of our Random Survival Forest (RSF) model, we generated time-dependent receiver operating characteristic (ROC) curves and computed the cumulative dynamic area under the curve (AUC) at multiple time points (at 10% of the time scale for each cohort, then 20%, 30%, 40%, 50%, 60%, 70%, 80% and 90%). The time-dependent AUC reflects the model’s ability to discriminate between individuals who experience an event (e.g., death) by a given time versus those who do not, accounting for censoring. An AUC close to 1 indicates strong predictive performance, while 0.5 corresponds to random chance. Additionally, we plotted the Kaplan-Meier survival curve based on observed outcomes in the test cohort, alongside the average predicted survival curve derived from the RSF model. This comparison provides a visual assessment of how closely the model’s predicted survival aligns with actual patient outcomes over time.

## Acknowledgements

The authors thank Yan Souza and Diego Chowell for helpful discussions. The results presented here are in whole or part based upon data generated by the TCGA Research Network: https://www.cancer.gov/tcga. A preprint version of this article has been peer-reviewed and recommended by PCI StatML (https://doi.org/10.24072/pci.statml.100105).

## Data Availability

All data are openly available on the TCGA website (data release 43, GRCh38/hg38). https://portal.gdc.cancer.gov/analysis_page?app=

Code is available on Github. https://github.com/gmestrallet/brcatcga

## Supplementary data

**Supplementary Table 1.**
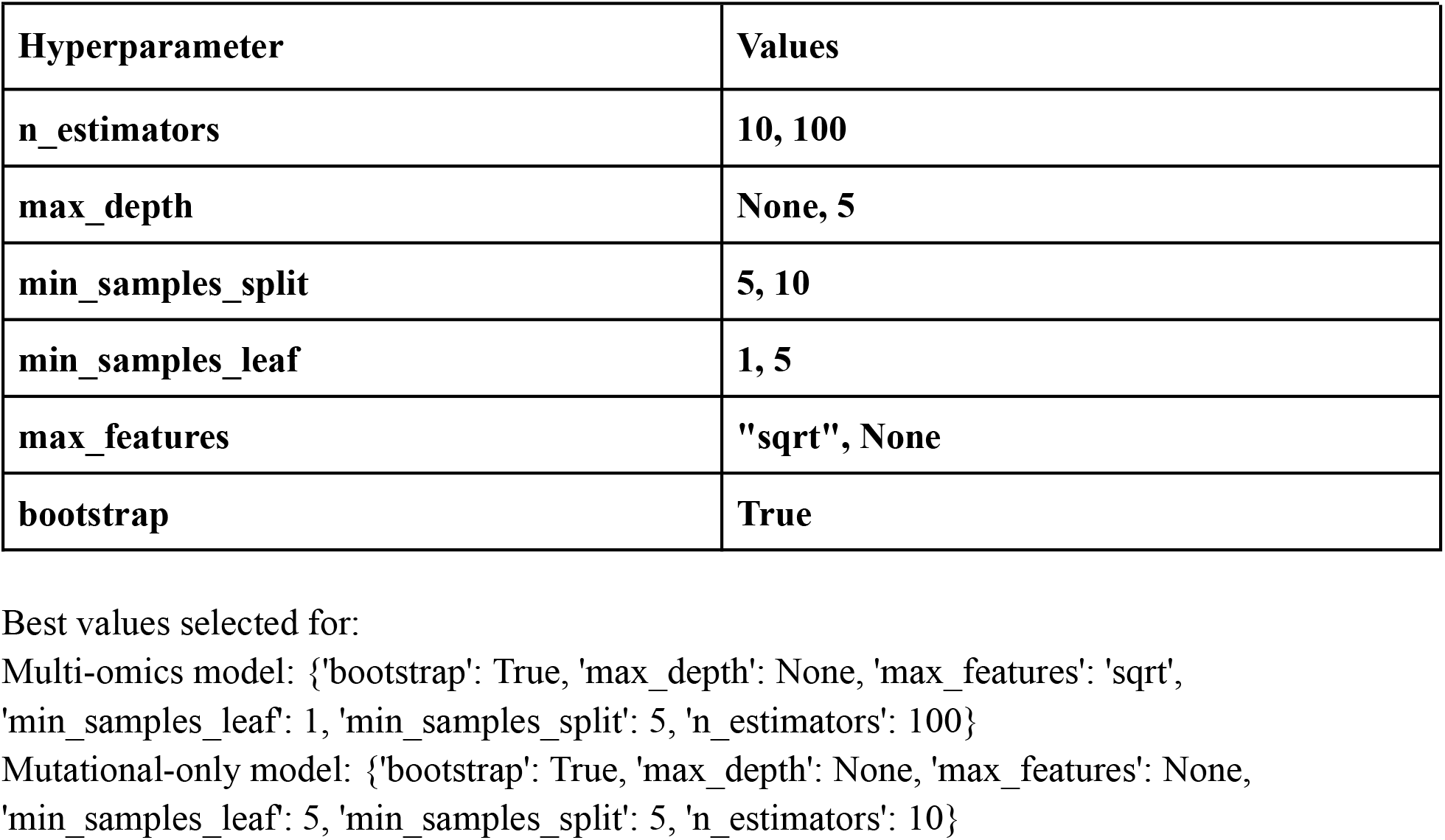
Hyperparameter search space.

| Hyperparameter | Values |
| --- | --- |
| **n_estimators** | **10, 100** |
| **max_depth** | **None, 5** |
| **min_samples_split** | **5, 10** |
| **min_samples_leaf** | **1, 5** |
| **max_features** | **"sqrt", None** |
| **bootstrap** | **True** |

Best values selected for:
Multi-omics model: {'bootstrap': True, 'max\_depth': None, 'max\_features': 'sqrt', 'min\_samples\_leaf': 1, 'min\_samples\_split': 5, 'n\_estimators': 100}
Mutational-only model: {'bootstrap': True, 'max\_depth': None, 'max\_features': None, 'min\_samples\_leaf': 5, 'min\_samples\_split': 5, 'n\_estimators': 10}

**Supplementary Figure 1.**
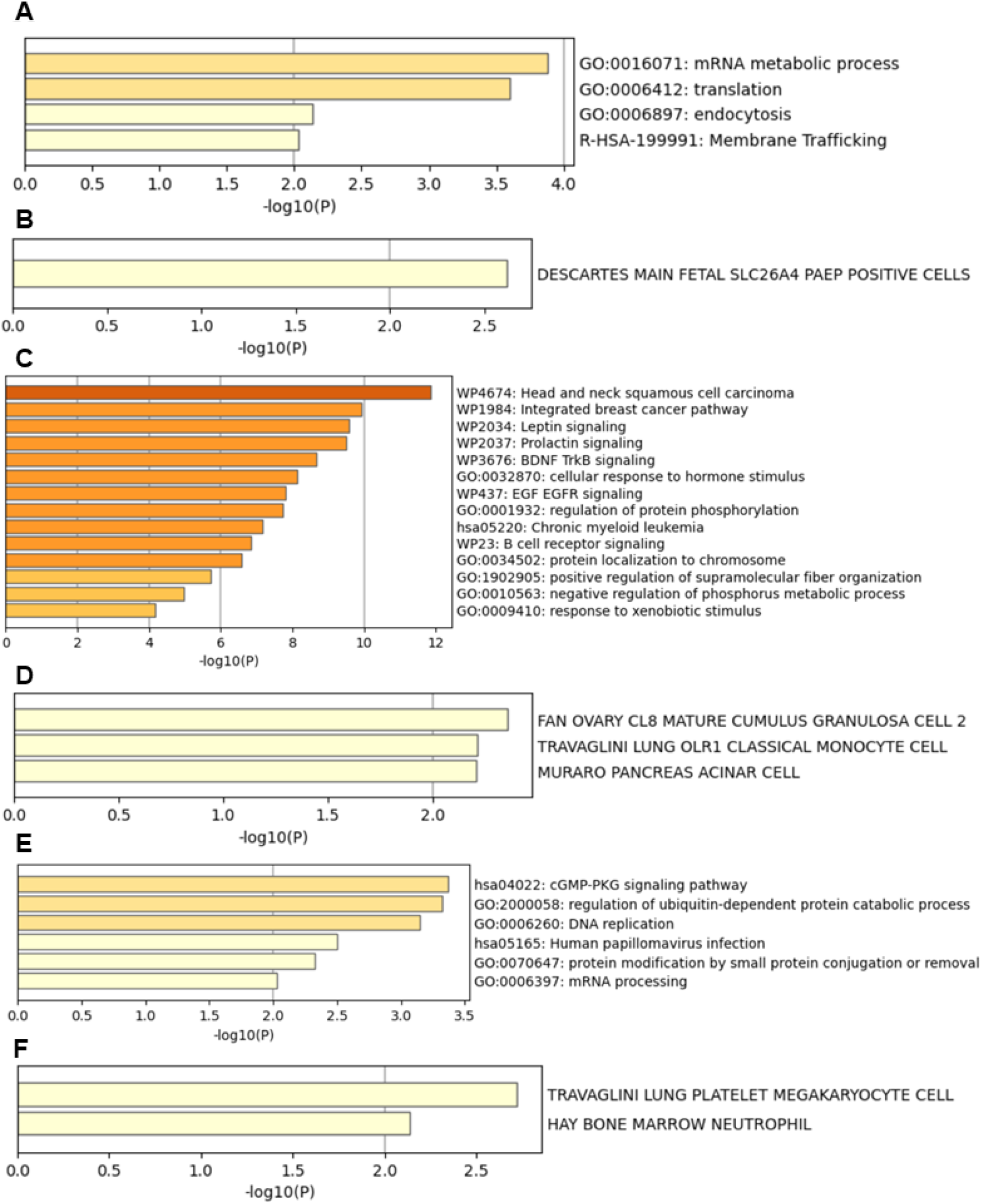
Breast cancer patient multi-omics and clinical features associated with a shorter survival and gene enrichment analysis. Gene enrichment (**A/C/E**) and cell type signature (**B/D/F**) analysis with these top/bottom genes ( **A/B** from the RNA dataset of Figure 1 C, **C/D** from the methylation dataset of Figure 1 D and **E/F** from the protein dataset of Figure 1 E) was performed using metascape (Y. Zhou et al., 2019).

**Supplementary Figure 2.**
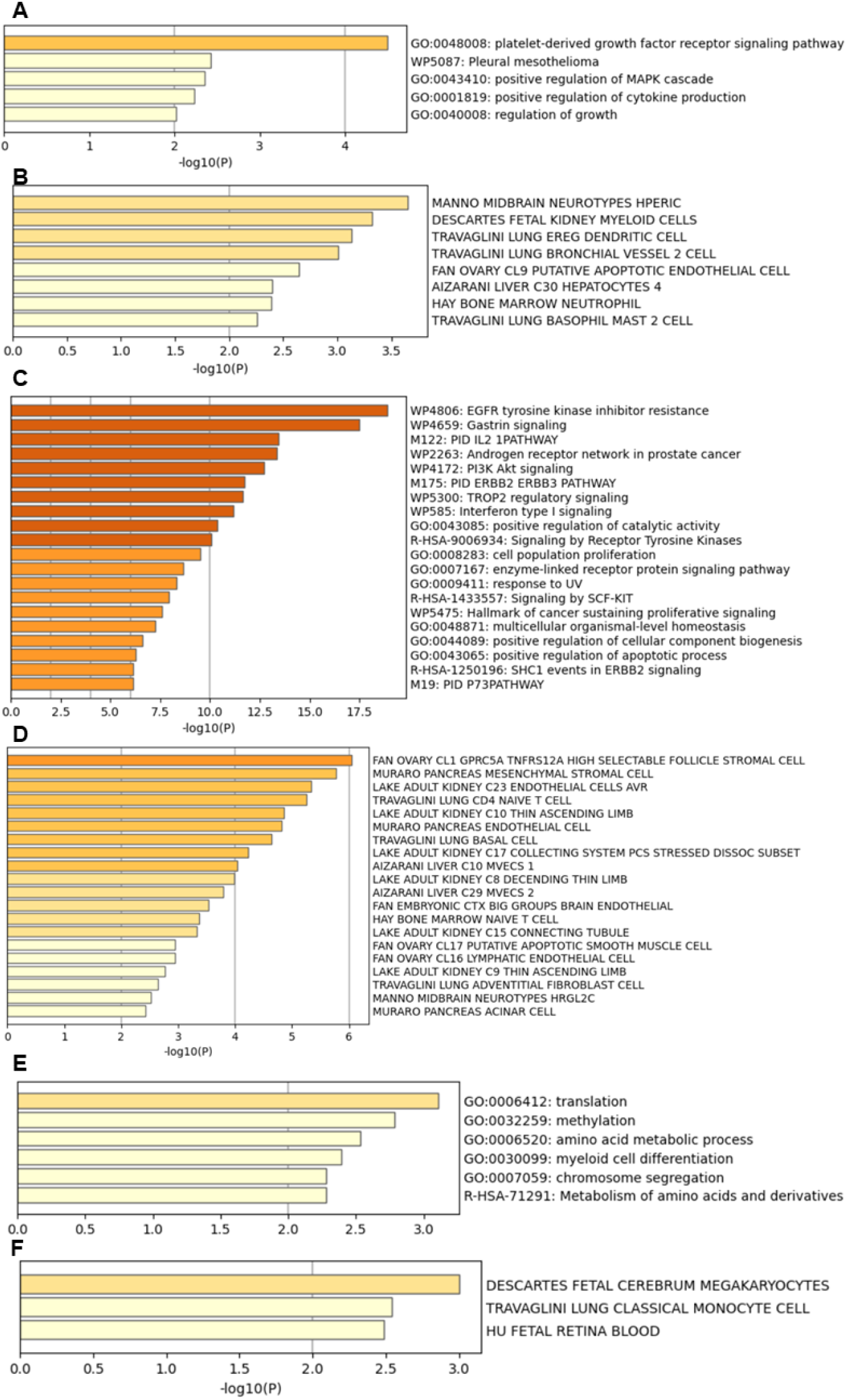
Breast cancer patient multi-omics and clinical features associated with a longer survival and gene enrichment analysis. Gene enrichment (**A/C/E**) and cell type signature (**B/D/F**) analysis with these top/bottom genes ( **A/B** from the RNA dataset of Figure 1 C, **C/D** from the methylation dataset of Figure 1 D and **E/F** from the protein dataset of Figure 1 E) was performed using metascape (Y. Zhou et al., 2019).

**Supplementary Figure 3.**
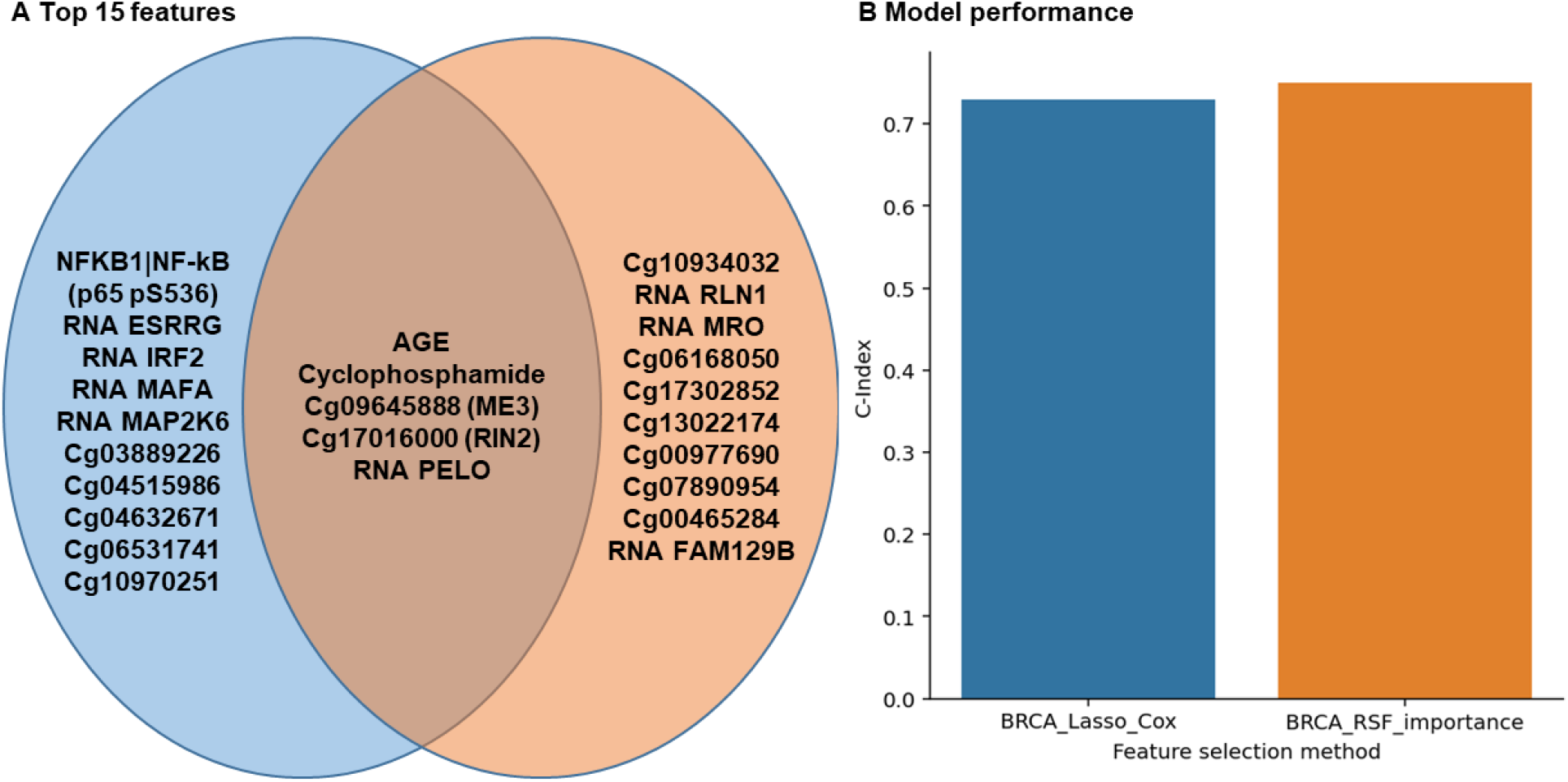
Comparison of model performance according to the feature selection method. **A**. Feature selection using Random Survival Forest permutation importance (orange) or Lasso-Cox (blue). N=802 breast cancer female patients. **B**. Model performance (C-Index) according to the feature selection method (RSFpermutation importance vs Lasso-Cox).

## Institutional Review Board Statement

Not applicable.

## Informed Consent Statement

Not applicable.

